# Role of transporters and enzymes in metabolism and distribution of 4 chlorokynurenine and metabolites

**DOI:** 10.1101/2023.07.14.548888

**Authors:** Waseema Patel, Ravi G. Shankar, Mark A. Smith, H. Ralph Snodgrass, Munir Pirmohamed, Andrea Jorgensen, Ana Alfirevic, David Dickens

## Abstract

4-chlorokynurenine (4-Cl-KYN) is in clinical development for potential CNS indications. We have sought to further understand the distribution and metabolism of 4-Cl-KYN as this information might provide a strategy to enhance the clinical development of this drug. We used excretion studies in rats, *in vitro* transporter assays and pharmacogenetic analysis of clinical trial data to determine how 4-Cl-KYN and metabolites are distributed. Our data indicated that a novel acetylated metabolite (N-acetyl-4-Cl-KYN) did not affect the uptake of 4-Cl-KYN across the blood-brain barrier via LAT1. 4-Cl-KYN and metabolites were found to be renally excreted in rodents. In addition, we found that N-acetyl-4-Cl-KYN inhibited renal and hepatic transporters involved in excretion. Thus, this metabolite had the potential to limit the excretion of a range of compounds. Our pharmacogenetic analysis found that a SNP in N-acetyltransferase 8 (NAT8, rs13538) was linked to levels of N-acetyl-4-Cl-KYN relative to 4-Cl-KYN found in the plasma and that a SNP in SLC7A5 (rs28582913) was associated with the plasma levels of the active metabolite, 7-Cl-KYNA. Thus, we have a pharmacogenetics-based association for plasma drug level that could aid in the drug development of 4-Cl-KYN and have investigated the interaction of a novel metabolite with drug transporters.

## Introduction

Major depressive disorder is an illness affecting millions of people worldwide, with approximately 3.8% of the population affected. The disorder is characterised by a myriad of symptoms including feelings of low mood, fatigue, insomnia, and anhedonia [1]. Despite the high incidence of the disorder, current treatment regimens are unable to treat up to one-third of patients. Furthermore, in patients who benefit, a process of trial and error may be required to determine the best dose and drug for them resulting in a lag time of weeks or even months between the initiation of treatment and improvement in symptoms [2].

Recent advances in intervention strategies for depression have focused on the glutamatergic system as a target for medicines development. Examples include 4-chlorokynurenine (4-Cl-KYN), esketamine and SLS-002 [1, 3–6]. Indeed, 4-Cl-KYN, a chlorinated form of kynurenine, a tryptophan derivative, has been investigated as an anti-depressant with potential for treatment for post-traumatic stress disorder [7, 8]. 4-Cl-KYN is thought to bring about anti-depressant effects via antagonism of the glycine_B_ site of the NMDA receptor (NMDAR). Specifically, 4-Cl-KYN is metabolised to the active metabolite, 7-chlorokynurenic acid (7-Cl-KYNA) within astrocytes in the CNS and this active compound then causes antagonism of the NMDAR. Pre-clinical data from rat studies have shown that 4-Cl-KYN can have rapid anti-depressant effects [4].

However, in spite of the promising pre-clinical data on 4-Cl-KYN as an anti-depressant, 4-Cl-KYN was unsuccessful in two phase 2 clinical trials, leading to a re-evaluation of its antidepressant properties in humans [9, 10]. The compound is still in clinical development in combination with probenecid (NCT05280054) to determine if this boosts the CNS concentration of the active metabolite in humans. This approach is based on a preclinical study in rodents that showed up to 885-fold higher brain extracellular fluid concentration of the active metabolite (7-Cl-KYNA) when probenecid was co-administered with 4-Cl-KYN [11]. Our last study comprehensively characterised the interaction of the 7-Cl-KYNA with numerous clinically relevant transporters [11]. Due to a number of positive preclinical studies, 4-Cl-KYN has a number of other potential indications that include as a therapy for L-DOPA induced dyskinesias, as an anticonvulsant and for use in neuropathic pain [12–15].

In this current study, we wanted to investigate the interaction of a newly discovered metabolite, N-acetyl-4-Cl-KYN (ac-4-Cl-KYN) with clinically relevant drug transporters. Furthermore, we sought to take advantage of the ELEVATE clinical cohort data (NCT03078322), and determine whether pharmacogenetic evaluation of our study cohort could provide novel insights into the relationship between drug concentrations and single nucleotide polymorphisms (SNPs). This pharmacogenetics approach may enable a personalised medicine strategy for 4-Cl-KYN drug development and dosing.

## Methods

### Cell culture

HEK 293 cells were grown at 37°C in a 5% CO_2_ humidified chamber. Cells were cultured in Dulbecco’s modified Eagles media (DMEM) supplemented with 10% foetal bovine serum (FBS).

### Stable cell line generation

HEK 293 cell lines stably expressing L type amino acid transporter (LAT) 1, and organic anion transporters 1, 2 and 3, were generated via stable transfection using Lipofectamine 2000 according to the manufacturer’s instructions, as described previously [11, 16]. In summary, transporters of interest (OAT1; NM_004790, OAT2 (NM_006672.3), OAT3; NM_001184732, or vector only) in the pcDNA3.1+/C-(K)DYK vector were introduced into HEK 293 cells, and cells expressing the vector were then selected for using G418 resistance. Selected cells were expanded, and single cell colonies were isolated. Cells from colonies with the highest expression were validated using qPCR and further expanded for use in experiments.

### Transient transfections

HEK 293 cells transiently expressing the transporters, human OAT1 (NM_004790) human OAT2 (NM_006672.3), human OAT3 (NM_001184732), OATP1B1 (NM_006446), rat (r) OAT1 (NM_017224.2), rOAT2 (NM_053537.2) and rOAT3 (NM_031332.1), or the matched empty vector (pcDNA3.1+/C-(K)DYK), were generated using Lipofectamine 2000 according to the manufacturer’s instructions. Briefly, HEK 293 cells were seeded at a density of 1 x 10^6^ cells/well the day before being transfected. Cells were then transfected with 2.5µg of DNA/well and experiments conducted 24 hrs after transfection.

### Drugs and radiochemicals

7-Cl-KYNA was obtained from Abcam and dissolved in a small volume of 1M HCl, the pH was then titrated to 7.4 in PBS. 4-Cl-KYN and ac-4-Cl-KYN were kind gifts from Vistagen Therapeutics, Inc. All other compounds were purchased from Sigma-Aldrich and solubilised in dimethyl sulfoxide according to manufacturer’s instructions.

[^14^C]-4-Cl-KYN (specific activity= 55.5 mCi/mmol) and [^14^C]-7-Cl-KYNA (specific activity= 77.7 mCi/mmol) were purchased from Moravek Inc. [^14^C]-tetraethylammonium specific activity= 3.5 mCi/mmol) and [^3^H]-estrone 3-sulfate specific activity= 49.19 Ci/mmol) were purchased from Perkin Elmer. [^3^H]-phenylalanine (specific activity= 100 Ci/mmol), [^3^H]-para-aminohippuric acid (specific activity= 40 Ci/mmol) and [^3^H]-cyclic guanosine monophosphate (specific activity= 25 Ci/mmol) were obtained from American Radiolabeled Chemical Inc. Finally, [^3^H]-L-DOPA (specific activity 4 Ci/mmol) was acquired from American Radiolabeled Chemicals, Inc.

### Trans-stimulation assays

HEK 293 cells stably expressing either LAT1 or matched empty vector were plated the day before an experiment; cells were 90% confluent at the time of the experiment. Culture media was first aspirated off the cells and cells were then washed with Hanks solution. Next, HEK 293 cells were exposed to radiolabelled substrate (L-DOPA) for 3 mins at 37°C following which, excess substrate was removed, and cells were again washed with Hanks solution. Cells were then exposed to unlabelled substrate for 3 mins at 37°C and excess substrate again removed. Finally, cells were washed with ice-cold Hanks solution. To determine intracellular radioactivity, cells were lysed with 5% sodium dodecyl sulphate (SDS) and scintillation fluid was added [11].

Data are presented as mean ± SD unless otherwise stated. Data from trans-stimulation assays were analyzed by one-way ANOVA. A p value <0.05 was considered statistically significant. Analyses were carried out on Graphpad Prism v8.

### Radiolabelled uptake assays

HEK 293 cells were plated either one or two days before an experiment, dependent on whether cells stably or transiently expressed the transporter of interest. Cell culture media was removed at the start of each experiment and cells were washed with Hanks balanced salt solution (25 mM HEPES, pH 7.4). Cells were then exposed to the radiolabelled compound of interest at a concentration of 0.15 µCi/ml for 3 mins at 37°C. Excess radiolabelled substrate was then removed, and cells were washed with ice cold Hanks to prevent further uptake. Intracellular uptake of the radiolabelled compound was determined by lysing cells with 5% SDS and adding scintillation fluid [11, 17].

Data are presented as mean ± SD unless otherwise stated. Data from uptake assays were analyzed by one-way ANOVA. A p-value <0.05 was considered statistically significant. Analyses were carried out on Graphpad Prism v8.

### Excretion mass balance and metabolite identification study

Animal experiments were conducted by Charles Rivers Laboratories (Ashland, USA) with the approval from their Institutional Animal Care and Use Committee. Food and water were made available to animals *ad libitum*. To determine the excretion of [^14^C]-4-Cl-KYN and/or metabolites, 3 male and 3 female Sprague-Dawley rats were orally dosed (gavage) with 500 mg/kg and 100 μCi/kg to determine routes of elimination and excretion mass balance. Following dosing, animals were placed into metabolism cages for separate collection of urine and faeces through 7 days. Following dosing, all excretion mass balance animals were placed in plastic metabolism cages for separate collection of urine and faeces. Samples were analysed for total radioactivity by liquid scintillation counting (LSC).

For the metabolite identification study, Sprague Dawley rats were dosed with a single oral dose of [^14^C]-4-Cl-KYN at 500 mg/kg and a target radioactivity of 250 μCi/kg. At 1 and 4 hours post dose, blood samples were collected from 1 animal/sex/time point. Selected plasma samples were profiled for metabolites using radiomatic-HPLC comprising a β-Ram model 4B radiomatic detector (IN/US Systems, Inc. Florida, USA) and a Zorbax SB-C18 column. Metabolites representing ≥5% of the total radioactivity (plasma) were analyzed by radio-HPLC/ mass spectrometry (MS)-MS to identify the radiolabeled metabolites.

### Clinical study cohort

A subset of study participants from the ELEVATE (ClinicalTrials.gov Identifier: NCT03078322) trial who provided informed written consent for pharmacogenetic analysis were used for our study. Participants were recruited from across the United States and were of mixed ethnic background, aged between 18 – 65 years. Trial participants had been previously diagnosed with major depressive disorder and were currently experiencing a depressive episode of at least 8 weeks in duration. For the trial, participants took a placebo or one oral dose of 4-Cl-KYN (1.44 g) in the morning after no/light breakfast in addition to an anti-depressant prescribed by their doctor for two weeks. A blood sample was taken from participants after taking an oral dose of 4-Cl-KYN.

The time between administration of 4-Cl-KYN and blood sample being taken was recorded. For each individual patient, at least one additional blood sample was taken on a separate occasion. 4-Cl-KYN, ac-4-Cl-KYN and 7-Cl-KYNA plasma concentrations were determined using HPLC with MS/MS detection by MicroConstants (San Diego, USA). In brief the method is applicable for measuring concentrations of 4-Cl-KYN, ac-4-Cl-KYN and 7-Cl-KYNA ranging from 50.0 to 50,000 ng/mL, 10.0 to 10,000 ng/mL and 2.00 to 2,000 ng/mL, respectively, using 40.0 μL of human plasma for extraction. The extracts were analyzed by HPLC using a Phenomenex Synergi MAX-RP column. The mobile phase was nebulized using heated nitrogen in electrospray positive ionization mode and the compounds were detected using MS/MS.

### Genotyping and imputation

Genotyping was performed by Genuity Science (Dublin, Ireland) using an Illumina Infinium global screening array. The tool gtc2vcf (Giulio Genovese, 2022, URL-https://github.com/freeseek/gtc2vcf) was used to convert intensity data files into VCF files for downstream analyses. Standard GWAS quality control was performed against the genotype data using PLINK 1.9 (https://zzz.bwh.harvard.edu/plink/cite.shtml) to filter out SNPs and individuals. The steps included removing samples with >1% missingness, duplicates or related samples, outlying homozygosity values, sex discordance and removal of ancestry outliers. Variant level checks removed SNPs with lower minor allele frequency and statistically significant Hardy-Weinberg Equilibrium values. The Michigan Imputation Server (https://github.com/genepi/imputationserver) was used for genotype imputation, as some of the SNPs to be investigated were not directly genotyped. The 1000 Genomes Phase 3 version 5 reference panel was used for the imputation. A total of 98 patient samples passed quality control analyses and were taken forward for analysis.

Due to the sample size, a targeted analysis based on a candidate gene approach for key SNPs was performed. Selected SNPs, from associations in metabolome-wide association studies (MWAS) that have been linked to the plasma level of the non-chlorinated derivatives of the kynurenine pathway, are summarized in Table 1. These SNPs were evaluated in a targeted analysis for association testing of SNPs and plasma concentrations.

**Table 1.** Association testing plan.

| **Individual Drug** | **Gene** | **Genotyping** | **Reason for testing** |
| --- | --- | --- | --- |
| 4-Cl-KYN | SLC7A5<br>(rs28582913) | Imputation | Shin et al 2014 & Long et al 2017 |
| 7-Cl-KYNA | KMO<br>(rs61825638) | Imputation | Shin et al 2014 & Long et al 2017 |
|  | SLC7A5<br>(rs28582913) | Imputation | Shin et al 2014, Long et al 2017 & Patel et al 2021 |
| ac-4-Cl-KYN | NAT8 (rs13538) | Direct | Yin et al 2022 |

**Table 1. Association testing plan.**
| Drug Ratios | Gene | Genotyping | Reason for testing |
| --- | --- | --- | --- |
| 7-Cl-KYNA/4-Cl-KYN | KMO<br>(rs61825638) | Imputation | Shin et al 2014 & Long et al 2017 |
|  | SLC7A5<br>(rs28582913) | Imputation | Shin et al 2014, Long et al 2017 & Patel et al 2021 |
| ac-4-Cl-KYN/4-Cl-KYN | NAT8 (rs13538) | Direct | Yin et al 2022 |

### Analysis for association between SNPs and plasma concentrations

The outcome variables we investigated were the plasma concentrations of 4-Cl-KYN, 7-Cl-KYNA, ac-4-Cl-KYN, and the ratios of 7-Cl-KYNA:4-Cl-KYN and ac-4-Cl-KYN:4-Cl-KYN. We used a regression modelling framework like that employed by researchers investigating clozapine and its metabolites [18]. Briefly, we tested for association between these outcomes and SNPs of interest (Table 1) using a linear fixed effects model and a random effect at subject level. Confounding variables such as the time difference between drug being taken and blood being collected, age, ethnicity, and visit, were considered univariately. The model used to test for association between plasma concentrations and SNPs then included the random effects at subject level and the variables found significant univariately, as well as the first two principal components of population structure. Where the distribution of plasma concentrations was non-Gaussian, the data were transformed using either the log_10_ or square-root transformation, as appropriate. Due to the 7 tests for association undertaken, we set our significance threshold to 0.05/7=0.0071 to reflect a Bonferroni correction for multiple testing.

### In silico docking

The AlphaFold predictive structure of human NAT8 (Q9UHE5-F1 v4) was used as a template for *in silico* docking. Identification of a putative docking pocket was determined with AMDock (ver 1.5.2) software [19] in an unbiased manner. In brief, 4-Cl-KYN and acetyl-CoA were docked with NAT8 using AutoDock Vina [20] within AMDock software.

The docking pose with the highest energy coefficient was used for generation of images produced in PyMOL (ver 2.52).

## Results

### N-acetyl-4-chlorokynurenine does not inhibit or interact with LAT1 transporter

Plasma samples from Sprague Dawley rats dosed with [^14^C]-4-Cl-KYN were utilized for metabolite analysis to investigate the pharmacology of 4-Cl-KYN. To determine the number and relative concentrations of radiolabeled compounds, the plasma samples were profiled using HPLC with radiodetection and mass spectrometry. The primary component present in plasma was the parent 4-Cl-KYN, representing approximately 53 percent relative observed intensity (% ROI) (Fig. 1A). The second largest component was M2, an unknown metabolite, eluting at 26 minutes, representing about 41% ROI. The third largest component from plasma was M1, representing about 6% ROI and is the known active metabolite, 7-Cl-KYNA.

**Figure 1.**
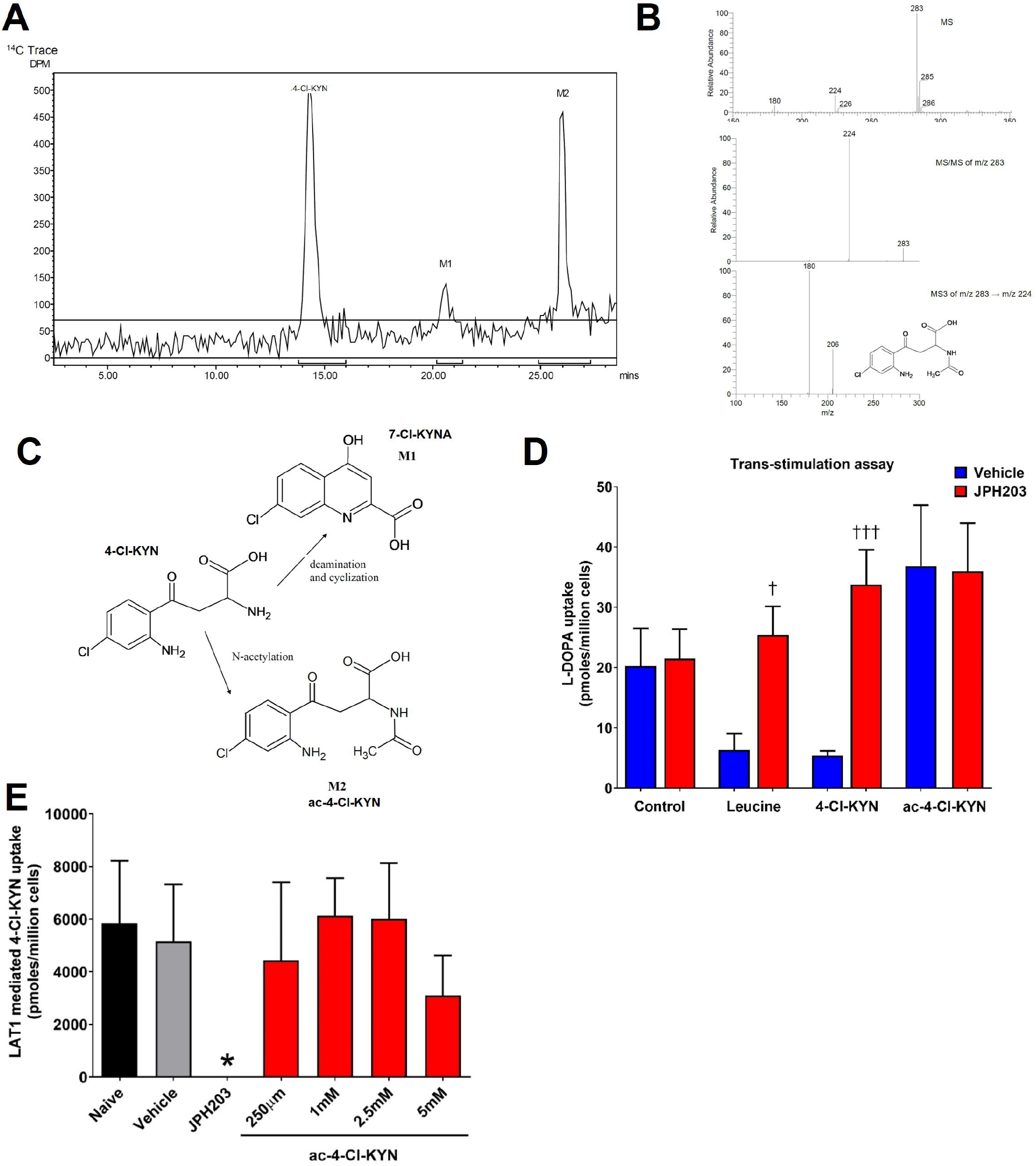
LAT1 transport is not affected by the novel metabolite, N-acetyl-4-chlorokynurenine. **(A)** Representative HPLC radiochromatogram of plasma from Sprague Dawley rat at a 1-hour time point following oral administration of [^14^C]-4-CI-KYN. **(B)** Mass spectrometry analysis of novel metabolite peak (M2) from rat plasma to ID metabolite as a N­ acetylated derivative of 4-CI-KYN. **(C)** Schematic for the metabolism of 4-CI-KYN to active metabolite (7-CI-KYNA, M1) or acetylation to generate ac-4-CI-KYN (M2). **(D)** HEK 293-LAT1 cells were preloaded with [3H]-levodopa (L-DOPA; 1 µM) and then exposed to 1 mM leucine, 4-chlorokynurenine (4-CI-KYN) or N-acetyl-4-chlorokynurenine (ac-4-CI-KYN) in the presence or absence of the inhibitor, JPH203 (10 µM). **(E)** Uptake of [^14^C]-4-CI-KYN in HEK 293 cells under conditions indicated. LAT1 mediated uptake of 4-CI­ KYN was determined by subtracting the uptake in HEK 293 control cells from the uptake in HEK 293-LAT1. Cells were exposed 250 µM 4-CI-KYN in the presence of the concentrations of ac-4-CI-KYN indicated. Data are mean ± SD (n=3). tp<0.05, tttp<0.001 compared to matched vehicle treated cells. *p<0.05 compared to na’i’ve cells.

Mass spectrometry was utilised to identify the M2 peak. The (-)ESI MS spectrum has a (M-H)-ion at m/z 283 with a 37Cl isotope peak at m/z 285 (Fig. 1B). The molecular ion is consistent with 4-Cl-KYN that has been acetylated. The MS/MS spectrum of m/z 283 has a single fragment ion at m/z 224 from the loss of NH2COCH3 (Fig. 1B). The MS3 spectrum of m/z 283 → m/z 224 has fragment ions at m/z 206 from the loss of H2O and at m/z 180 from the loss of CO2 (Fig. 1B). The mass spectrometry data are consistent with a structure of 4-Cl-KYN that has undergone acetylation at the alpha amino position.

To summarise, the metabolic pathway for 4-Cl-KYN involves either N-acetylation at the alpha amino position to form M2 (N-acetyl-4-Cl-KYN) or 4-Cl-KYN undergoes cyclization with the loss of NH3 to form M1 (7-Cl-KYNA) (Fig. 1C).

We then the effect of N-acetyl-4-Cl-KYN (ac-4-Cl-KYN) on transporters expressed in the brain and elsewhere. Our previous study investigated the interaction of the active metabolite, 7-Cl-KYNA, with several transporters [11]. We started by investigating whether ac-4-Cl-KYN could interact with LAT1 (SLC7A5) using a trans-stimulation assay as this transporter is instrumental in the uptake of 4-Cl-KYN across the BBB. Thus, we pre-loaded HEK 293 LAT1 cells with a known radiolabelled substrate of LAT1 (L-DOPA), washed out excess substrate and then exposed cells to potential substrates for LAT1. We found that known substrates of LAT1 (such as leucine and 4-Cl-KYN) caused a reduction in intracellular L-DOPA accumulation due to the antiporter mechanism of LAT1 extruding L-DOPA for uptake of leucine or 4-Cl-KYN. The LAT1 inhibitor, JPH203, prevented the leucine and 4-Cl-KYN induced reduction in intracellular L-DOPA. ac-4-Cl-KYN, however, had no effect on intracellular L-DOPA accumulation, suggesting that it is not a substrate for LAT1 (Fig. 1D).

We next tested whether ac-4-Cl-KYN could affect 4-Cl-KYN uptake via LAT1 by acting as an inhibitor by measuring the uptake of 4-Cl-KYN in HEK 293-LAT1 and matched control cells in the presence of a series of concentrations of ac-4-Cl-KYN. We found that the LAT1 inhibitor JPH203 caused a reduction in the LAT1 mediated uptake of 4-Cl-KYN, but none of the concentrations of ac-4-Cl-KYN tested had any effect (Fig. 1E). This data indicated that the presence of ac-4-Cl-KYN does not interfere with LAT1 mediated uptake of 4-Cl-KYN.

### 4-chlorokynurenine and metabolites are excreted via urine

As acetylated metabolites can be subject to renal excretion and as the active metabolite of 4-Cl-KYN (7-Cl-KYNA) is a substrate of renal transporters [11], we wanted to further understand the distribution of 4-Cl-KYN and/or its metabolites. To investigate this, an *in vivo* excretion mass balance study was performed in Sprague Dawley rats following a single oral dose of [^14^C]-4-Cl-KYN. We found that 4-Cl-KYN and metabolites are primarily excreted via urine, evidenced by 76% [^14^C]-4-Cl-KYN related radioactivity recovered in urine (Fig. 2A). Radioactivity recovered in faeces was 14.9%. In both cases, no differences were observed in excretion of the compound between males and females.

**Figure 2.**
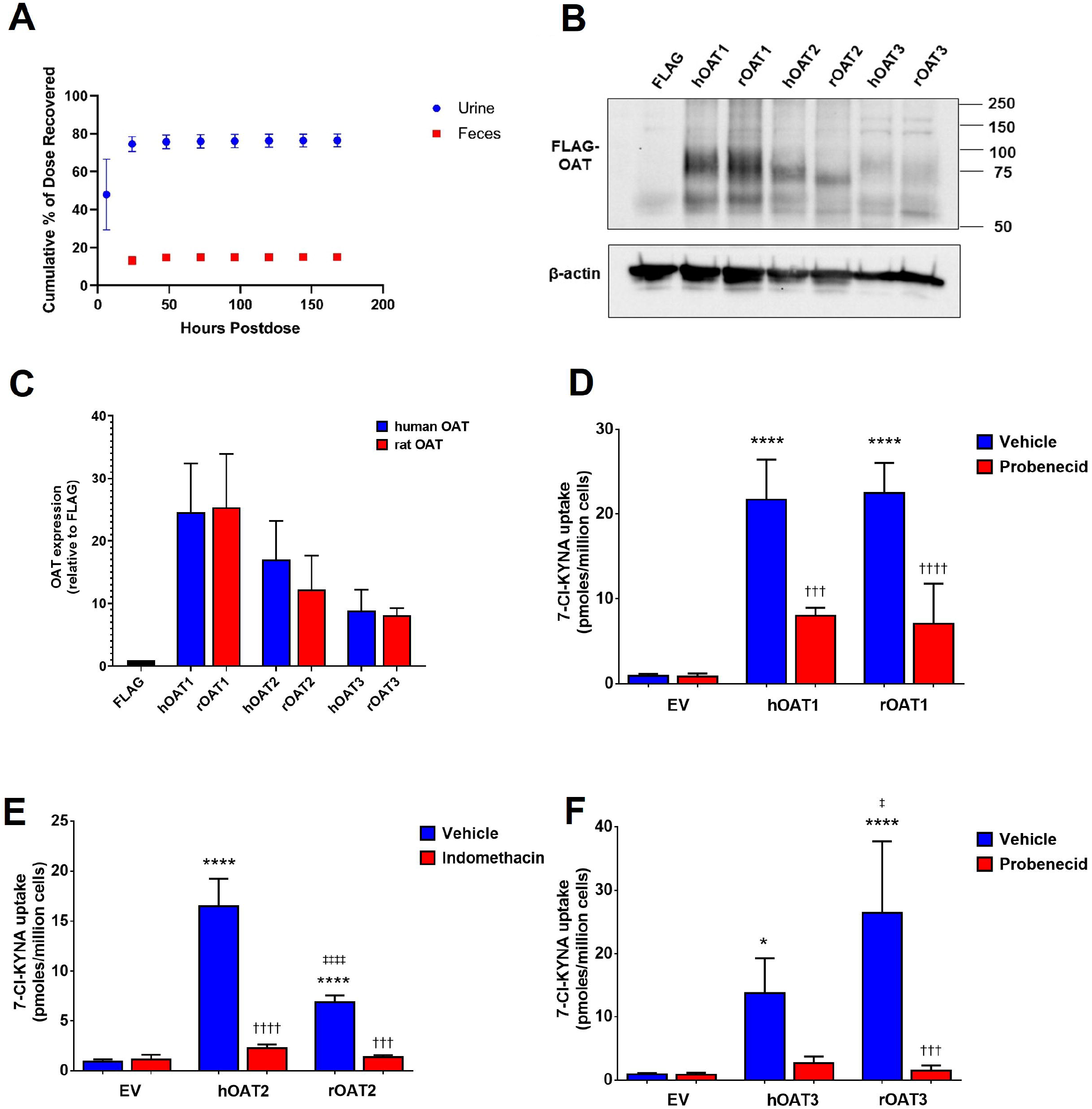
Differences in the uptake of 7-Cl-KYNA between human and rat organic anion transporters. **(A)** Routes of elimination and excretion mass balance of [^14^C]-4-CI-KYN derived radioactivity in Sprague Dawley rats following a single oral dose. Data are mean ± SD (n=6). **(B)** Representative western blot images showing the expression of human and rat FLAG-OATs under conditions indicated. **(C)** Densiometric quantification of the expression of OATs under conditions indicated. Densiometric quantification for OATs was carried out using the band at the expected size of 75 kDa. Changes in the expression of OATs were normalised to-actin and are shown relative to OAT expression in control cells (FLAG+HA). **(D-F)** Uptake of [^14^C]-7-chlorokynurenic acid (7-CI­ KYNA) under conditions indicated in control HEK 293 cells and HEK 293 cells transfected with human (h) or rat (r) organic anion transporters (OAT) **(D) 1 (E)** 2 or **(F)** 3. Cells were exposed 2 µM 7-CI-KYNA, 100 µM indomethacin or 1 mM probenecid. Data are mean ± SD (n=3). **p<0.01, ****p<0.0001 compared to empty vector (EV), tp<0.05, ttp<0.01, tttp<0.001, ttttp<0.0001

### Differences in the uptake of 7-Cl-KYNA between human and rat organic anion transporters

Species differences have been reported for OAT transporters, for example human versus rat OATs for the transport of kynurenic acid [21]. We next tested whether there were any species differences in OAT activity towards 7-Cl-KYNA transport. We used transient transfection to introduce human and rat homologs of OAT transporters 1,2 and 3 (SLC22A6, SLC22A7 & SLC22A8) into HEK 293 cells. Western blot studies validated the expression of the different OAT transporters and showed that there were no differences in the levels of expression of the different OAT transporters between the human and rat homologs (Fig. 2B+C).

We next sought to compare the uptake of 7-Cl-KYNA via the human and rat OATs. On average, we found uptake of 7-Cl-KYNA via hOAT1 (21.7 ± 4.8 pmoles/million cells) and rOAT1 (22.5 ± 3.6 pmoles/million cells) to be similar and probenecid to have a similar effect on both transporters, 8.0 ± 1.0 pmoles/million cells and 7.1 ± 4.8 pmoles/million cells for hOAT1 and rOAT1, respectively (Fig.2D). Conversely, we found that hOAT2 (16.5 ± 2.8 pmoles/million cells) had a higher uptake of 7-Cl-KYNA than rOAT2 (6.9 ± 0.7 pmoles/million cells, Fig.2E, n=3, p<0.05). We also found differences in uptake between hOAT3 and rOAT3; hOAT3 (13.8 ± 5.6 pmoles/million cells) was found to have a lower uptake of 7-Cl-KYNA than rOAT3 (26.4 ± 11.3 pmoles/million cells, Fig.3F, n=3, p<0.05). Due to these species differences, we focus additional experiments on human data from *in vitro* or clinical data to further investigate the pharmacology of these compounds.

**Figure 3.**
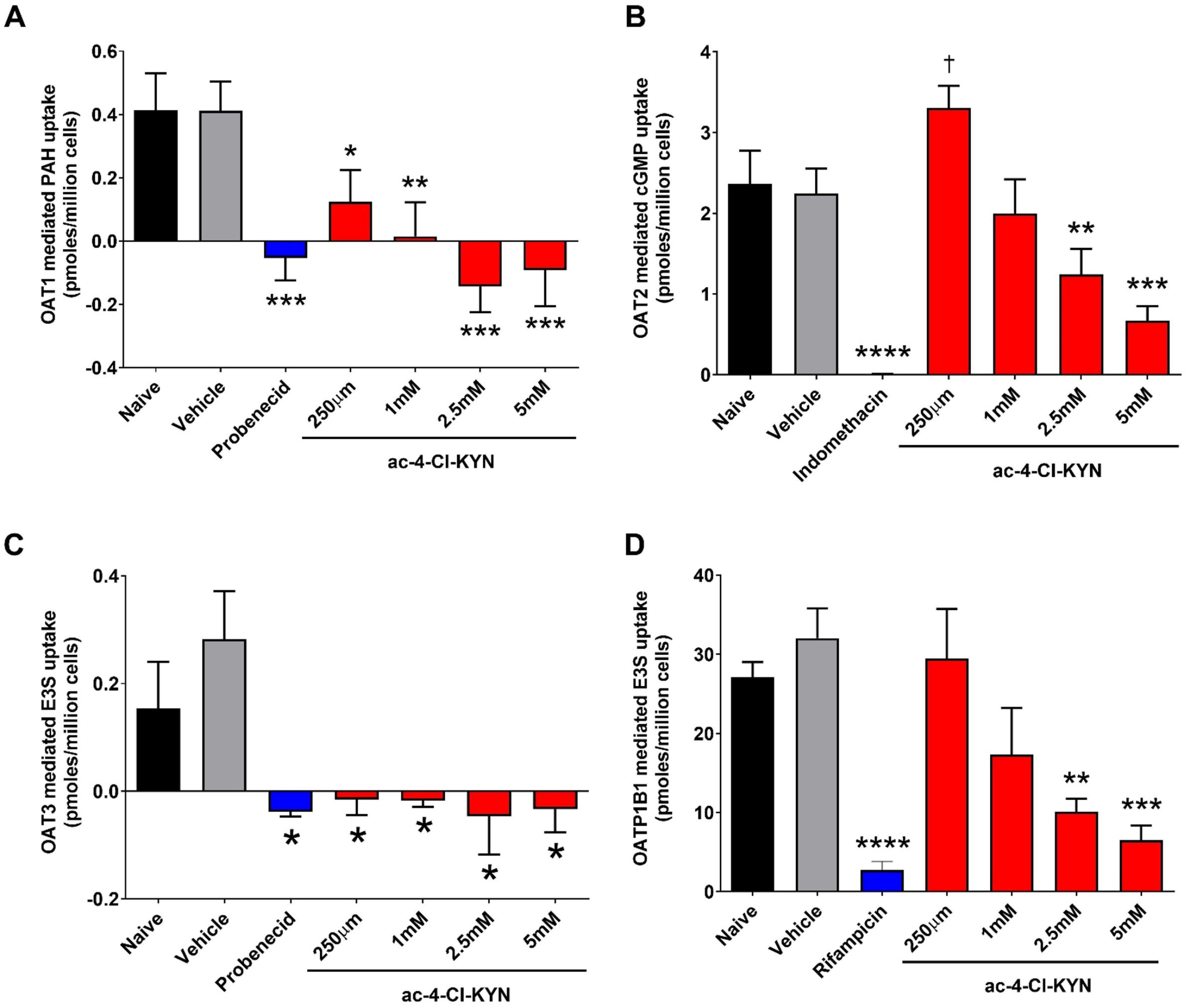
N-acetyl-4-chlorokynurenine inhibits renal and hepatic transporters. **(A)** Uptake of [3H]-para-aminohippuric acid (PAH; 1 µM) under conditions indicated. Organic anion transporter (OAT) 1 mediated uptake of PAH was determined by subtracting the uptake in HEK 293 control cells from the uptake in HEK 293-OAT1. **(B)** Uptake of [3H]-cyclic guanosine monophosphate (cGMP) under conditions indicated. OAT2 mediated uptake of cGMP was determined by subtracting the uptake in HEK 293 control cells from the uptake in HEK 293-OAT2. **(C)** Uptake of [3H]-estrone-3 sulphate (E3S; 100 nM) under conditions indicated. OAT3 mediated uptake of E3S was determined by subtracting the uptake in HEK 293 control cells from the uptake in HEK 293-OAT3. Data are mean ± SD (n=3). **(D)** Uptake of [3H]-estrone-3 sulphate (E3S; 100 nM) under conditions indicated. OATP1B1 mediated uptake of E3S was determined by subtracting the uptake in HEK 293 control cells from the uptake in HEK 293-OATP1B1. Cells were exposed to a model substrate in the presence or absence of different chemicals including a range of concentrations for ac-4-CI-KYN as indicated. Data are mean ± SD (n=3). *p<0.05, **p<0.01, ***p<0.001, ****p<0.0001

### N-acetyl-4-chlorokynurenine inhibits renal and hepatic transporters

We next determined whether the inactive metabolite of 4-Cl-KYN, N-acetyl-4-Cl-KYN, could affect other clinically relevant human drug transporters as this could be important in drug-drug interactions. We started by looking at the renally expressed transporter using our previously established stably expressing cell lines [11]. We found that the presence of N-acetyl-4-Cl-KYN caused a dramatic inhibition of OAT1 (*SLC22A6*) and OAT3 (*SLC22A8*) mediated uptake of the model substrates para-aminohippuric acid and estrone-3 sulphate, respectively (Fig. 3A & C).

We also studied the hepatic transporters OAT2 (*SLC22A7*) and OATP1B1 (*SLCO1B1*). We found that uptake of the OAT2 model substrate, cyclic guanosine monophosphate, was reduced in the presence of N-acetyl-4-Cl-KYN (Fig. 3B). Similarly, OATP1B1 mediated uptake was also reduced in the presence of N-acetyl-4-Cl-KYN (Fig. 3D). These data, together, indicate that the acetylated metabolite of 4-Cl-KYN is capable of inhibiting human transporters involved in excretion in both the kidneys and liver.

### Analyses of association between SNPs and individual drugs

Candidate SNPs, based on prior associations with kynurenine pathway metabolites (Table 1), were investigated for association with the plasma concentrations of 4-Cl-KYN and/or drug metabolites using a regression modelling framework. In univariate analyses with potential confounders, time from dosing was significantly associated with ac-4-Cl-KYN levels (p: 0.000007). The significant factor was adjusted for in the SNP association analyses, together with the first 2 principal components of ancestry.

No association was found between SLC7A5 rs28582913 and 4-Cl-KYN (p: 0.01) (Fig. 4A) or between KMO rs61825638 and 7-Cl-KYNA levels (p: 0.193) (Fig. 4B). An association was identified between LAT1 rs28582913 and 7-Cl-KYNA levels (p: 0.00667), and the SNP appeared to have a recessive effect, with two copies of the ‘C’ allele at rs28582913 leading to approximately 46% lower blood concentrations of 7-Cl-KYNA (Fig. 4C). Finally, no association was identified between the SNP in N-acetyltransferase 8 (NAT8; rs13538) and the concentration of ac-4-Cl-KYN (p: 0.07) (Fig. 4D & Table 2).

**Figure 4.**
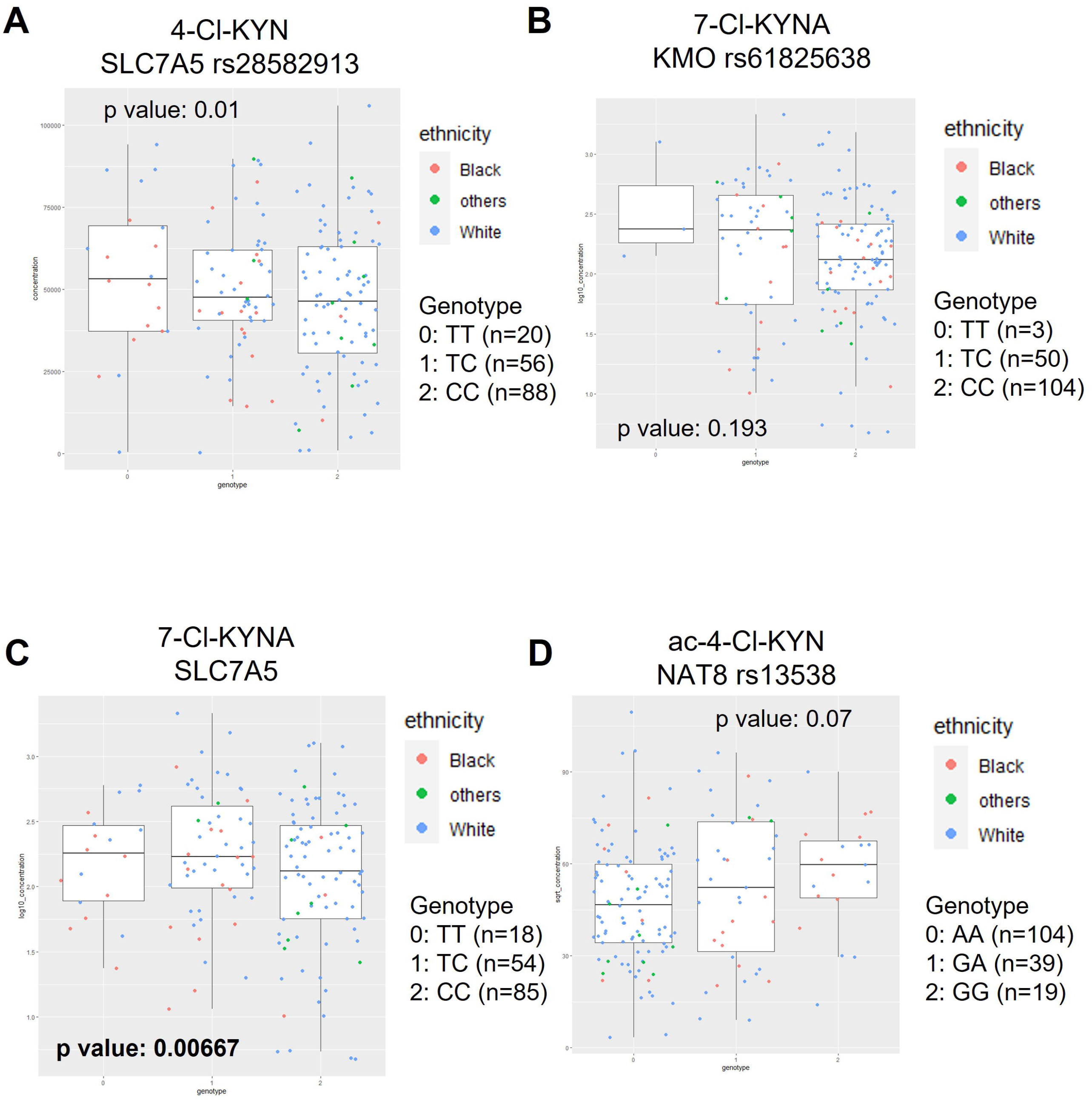
7-CI-KYNA levels are linked to a SLC7A5 SNP (rs28582913). **(A)** Effect of SLC7A5 SNP (rs28582913) on plasma level of 4-CI-KYN. **(B)** Effect of KMO SNP (rs61825638) on 7-CI­ KYNA plasma levels. **(C)** Association of 7-CI-KYNA plasma concentrations with SLC7A5 SNP (rs28582913). **(D)** Effect of NAT8 SNP (rs13538) on ac-4-CI-KYN plasma levels. Plasma levels of compounds of interest were determined from patients participating in the ELEVATE clinical trial. Significance threshold was p<0.05/7 with significant p values in bold. Self-reported ethnicity is shown with other ethnic background shortened to others.

**Table 2.**
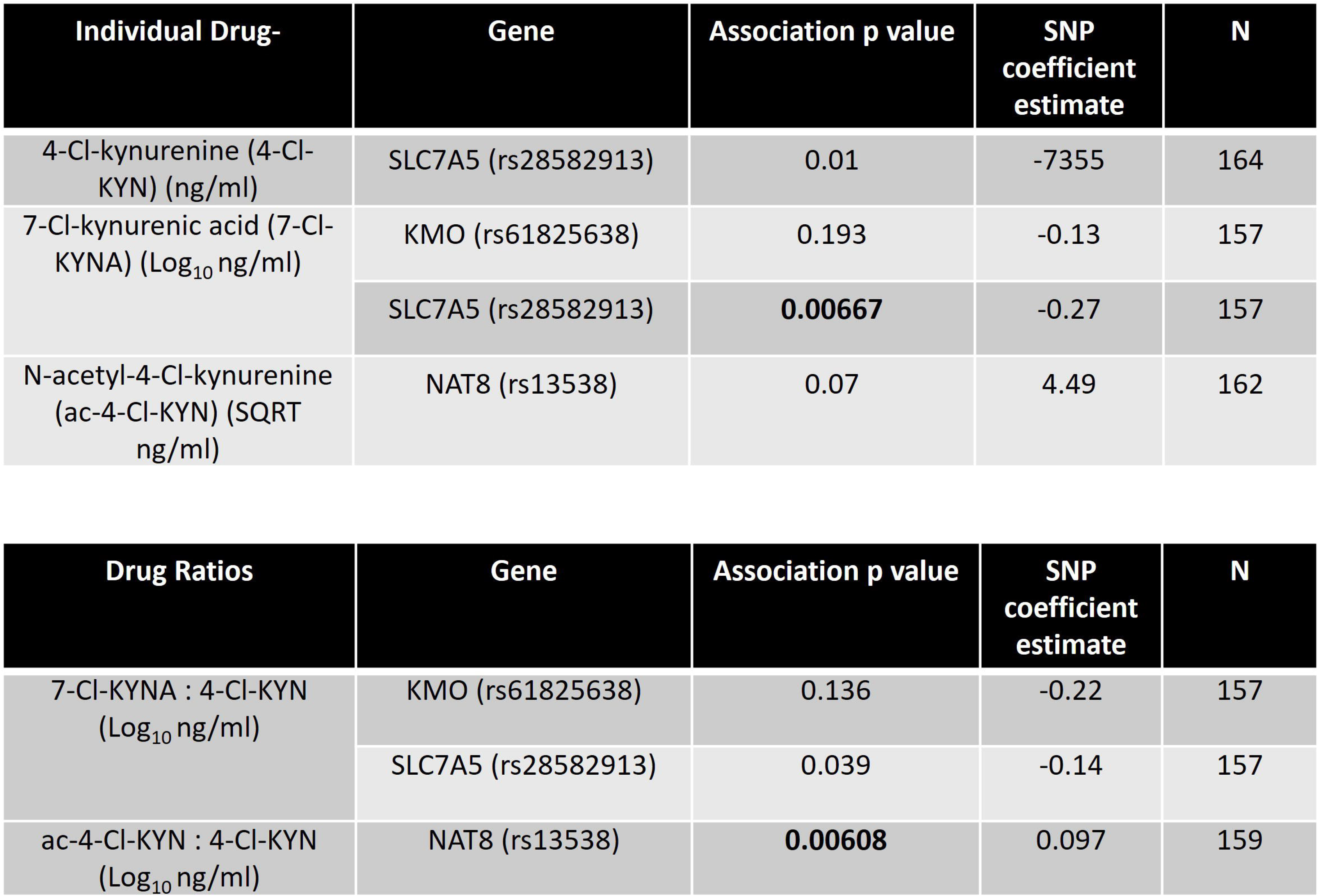
Summary of SNP and blood concentration association testing. Results of testing for association between SNPs and blood concentration of parent or metabolites of 4-CI-KYN using a linear mixed model regression modelling approach. Significance threshold was p<0.05/7. The following covariates were adjusted for; time from dosing, sex, and ethnicity.

| Individual Drug- | Gene | Association p value | SNP coefficient estimate | N |
| --- | --- | --- | --- | --- |
| 4-Cl-kynurenine (4-Cl-KYN) (ng/ml) | SLC7A5 (rs28582913) | 0.01 | -7355 | 164 |
| 7-Cl-kynurenic acid (7-Cl-KYNA) (Log<sub>10</sub> ng/ml) | KMO (rs61825638) | 0.193 | -0.13 | 157 |
|  | SLC7A5 (rs28582913) | **0.00667** | -0.27 | 157 |
| N-acetyl-4-Cl-kynurenine (ac-4-Cl-KYN) (SQRT ng/ml) | NAT8 (rs13538) | 0.07 | 4.49 | 162 |

**Table 2. Summary of SNP and blood concentration association testing.**
| Drug Ratios | Gene | Association p value | SNP coefficient estimate | N |
| --- | --- | --- | --- | --- |
| 7-Cl-KYNA : 4-Cl-KYN (Log<sub>10</sub> ng/ml) | KMO (rs61825638) | 0.136 | -0.22 | 157 |
|  | SLC7A5 (rs28582913) | 0.039 | -0.14 | 157 |
| ac-4-Cl-KYN : 4-Cl-KYN (Log<sub>10</sub> ng/ml) | NAT8 (rs13538) | **0.00608** | 0.097 | 159 |

### Association between metabolite:4-Cl-KYN ratios and SNPs

In univariate analyses with potential confounders, time from dosing was significantly associated with 7-Cl-KYNA:4-Cl-KYN (p: 0.025) and ac-4-Cl-KYN:4-Cl-KYN (p: 0.00000003) ratios. The significant factor was adjusted for in the SNP association analyses, together with the first 2 principal components of ancestry.

No association was found between KMO rs61825638 and 7-Cl-KYNA:4-Cl-KYN (p: 0.136), or between SLC7A5 rs28582913 and 7-Cl-KYNA:4-Cl-KYN (p: 0.039) (Fig. 5A&B). An association was identified between NAT8 rs13538 SNP and ac-4-Cl-KYN:4-Cl-KYN (p: 0.00608) (Fig 5C & Table 2). The results suggested that the SNP had an additive effect with a 1.86 fold difference in ratio seen between those with two ‘G’ alleles at SNP rs13538 compared to those with two ‘A’ alleles. This NAT8 SNP has been associated by eQTL analysis with gene expression and is also a non-synonymous SNP. We therefore investigate if this SNP has a direct functional role at the protein level by *in silico* modelling approaches.

**Figure 5.**
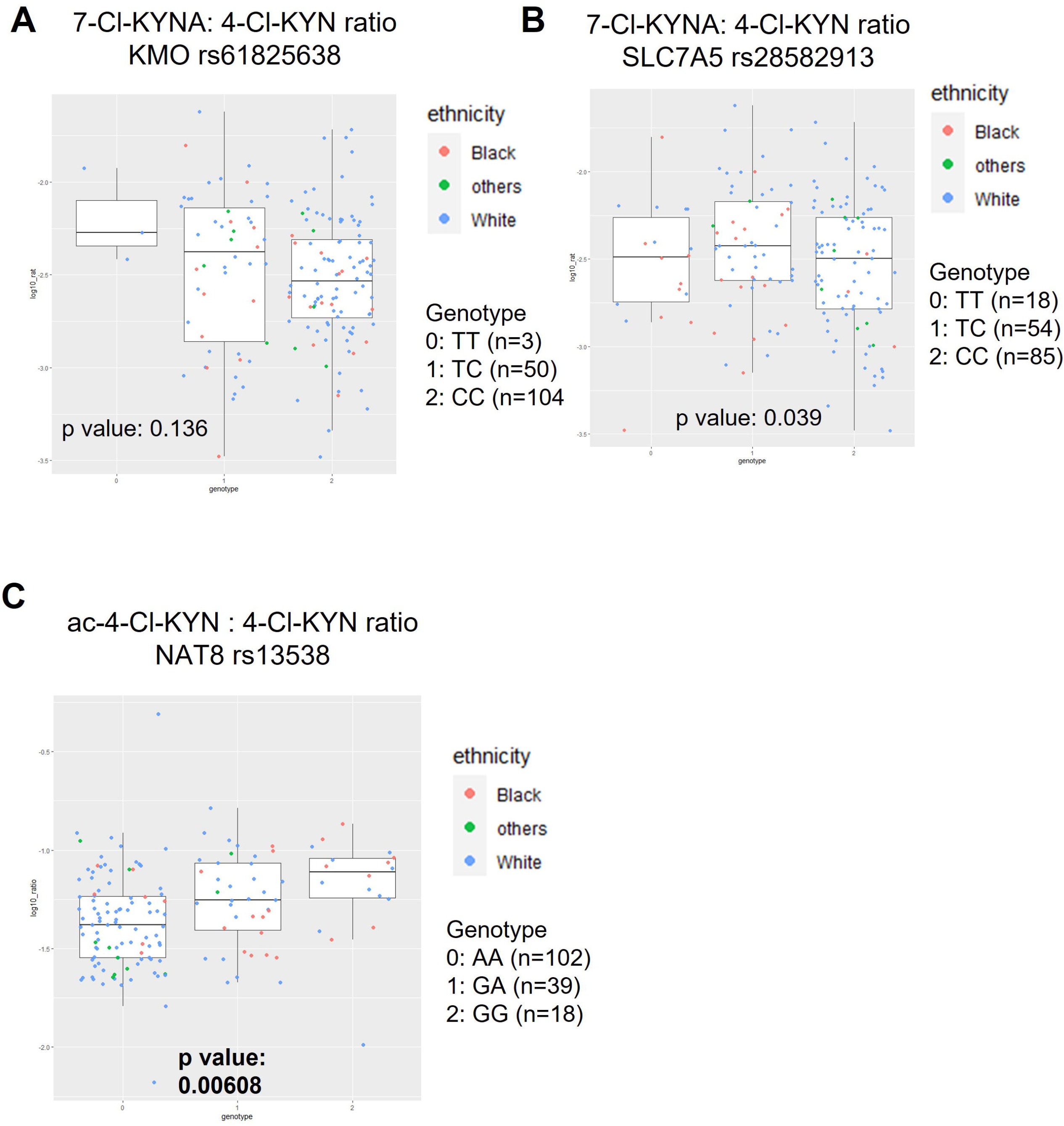
ac-4-CI-KYN:4-CI-KYN plasma ratio is associated with a NAT8 SNP (rs13538). **(A)** Effect of a KMO SNP (rs61825638) on plasma ratio of 7-CI-KYNA: 4-CI-KYN. **(B)** Effect of a SLC7A5 SNP (rs28582913) on plasma ratio of 7-CI-KYNA: 4-CI-KYN. **(B)** Effect of on 7-CI­ KYNA plasma levels. **(C)** Association of ac-4-CI-KYN: 7-CI-KYNA plasma ratio with a NAT8 SNP (rs13538). Plasma levels of compounds of interest were determined from patients participating in the ELEVATE clinical trial. Significance threshold was p<0.05/7 with significant p values in bold. Self-reported ethnicity is shown with other ethnic background shortened to others.

### Non-synonymous NAT8 SNP is located in the putative substrate binding pocket of NAT8

Taking advantage of an AlphaFold [22] model of human NAT8 (Fig.6A) we probe the predicted 3D structure of this putative acetyltransferase and location of the non-synonymous SNP (rs13538). The protein has a continuous pore (Fig.6 B&C) with two entrances. The channel is relatively large with a predicted solvent accessible pore of 2604 Å^3^. *In silico* docking of 4-Cl-KYN and acetyl-co-A (Fig.6 D&E) showed that the compounds utilise the breadth of the channel with the acetyl-co-A and 4-Cl-KYN in close proximity. The non-synonymous SNP (rs13538) changes Phe to Ser at amino acid 143 which is located at the putative interface of where the acetyl-co-A and 4-Cl-KYN interact at the binding pocket (Fig.6 F). This could suggest an important functional role of amino acid 143 in substrate turnover that is altered in individuals with Phe143Ser substitution.

**Figure 6.**
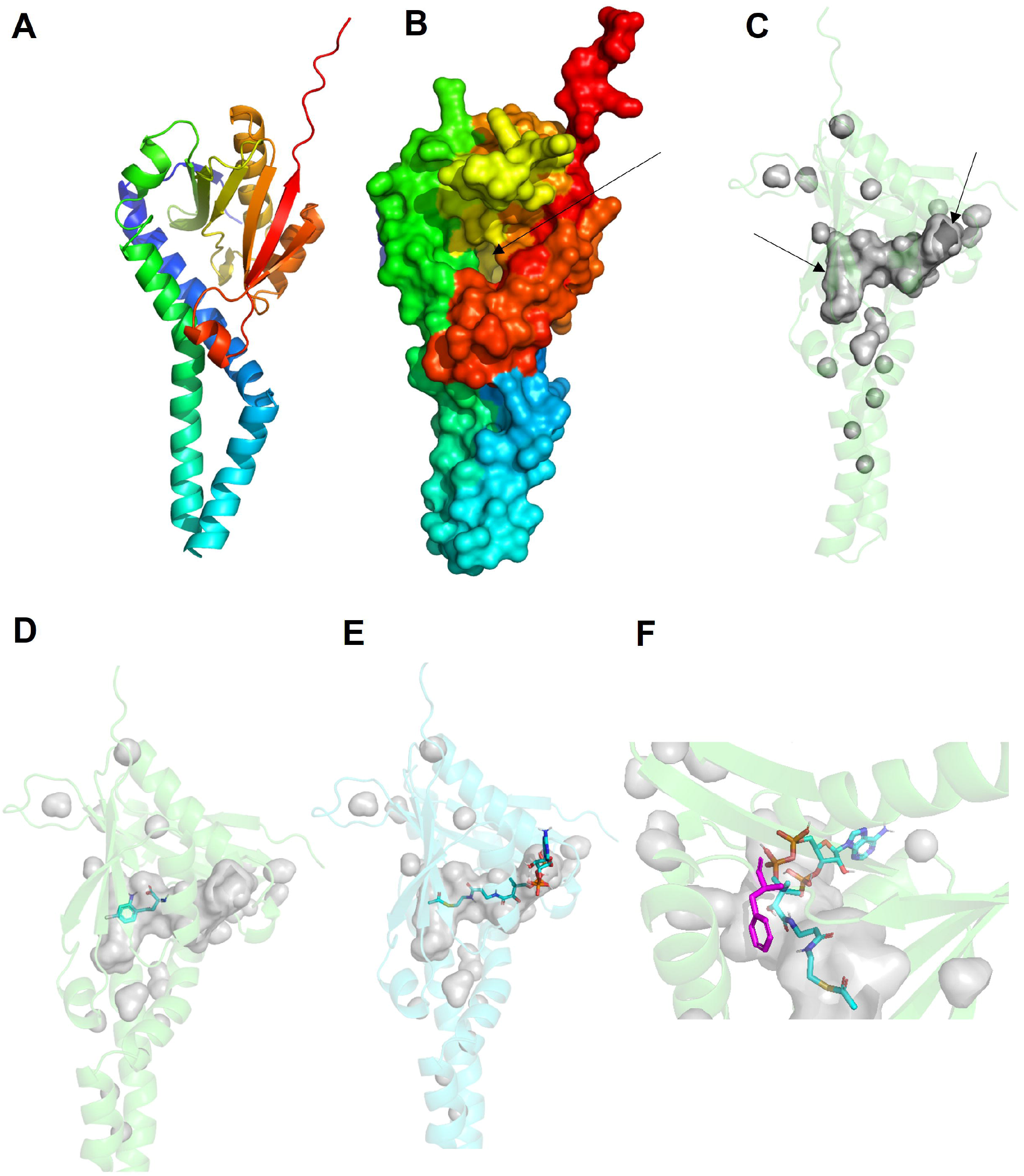
*In silica* docking of 4-CI-KYN & acetyl-CoA onto NAT8. **(A)** Overall AlphaFold predicted structure of NAT8 with secondary structures as cartoons. **(B)** Surface of NAT8 with focus on pore entrance marked by arrow. **(C)** Solvent accessible continuous pore of NAT8 in grey with arrows at either end of channel (active site). Docking of 4-CI-KYN **(D)** or acetyl Co-A **(E)** onto NAT8. **(F)** Location of non-synonymous SNP (Phe143Ser, rs13538) in purple on NAT8 with docked acetyl Co-A.

## Discussion

4-Cl-KYN is undergoing clinical development (NCT05280054) for a range of possible CNS related indications. To aid in the development of this compound we have identified and investigated a novel metabolite, N-acetyl-4-Cl-KYN. Once absorbed, 4-Cl-KYN is readily taken up across the BBB and converted to the active metabolite within astrocytes [23–25]. We found that N-acetyl-4-Cl-KYN had no effect on the transport activity of LAT1, indicating that this metabolite is unlikely to interfere with the uptake of 4-Cl-KYN across the BBB and thus will not affect the efficacy of the compound. This is similar to a finding by Nagamori *et al* that found that N-acetyl-leucine does not interact with LAT1 [26]. Our excretion studies found that 4-Cl-KYN and/or metabolites are excreted via urine. We found N-acetyl-4-Cl-KYN inhibited multiple transporters expressed at the basolateral membrane of renal proximal tubules and the sinusoidal membrane of hepatocytes. OATs and OATP1B1 are important for the clearance of compounds from the plasma into the urine. N-acetyl-leucine is also a substrate of OAT1 and OAT3 [26]. A recent MWAS approach has identified SLC17A4 as a putative N-acetyl kynurenine transporter as a SNP located near this gene was associated with N-acetyl kynurenine levels in the blood [27]. SLC17A4 has been shown by both functional and genetic experiments to be a thyroid hormone transporter [28, 29]. Drug-drug interaction potential and direct transport studies could be investigated in further work.

We investigated the potential of a pharmacogenetics approach that could be used to stratify patients. In a recent review article on psychiatry pharmacogenetics, Pardinas *et al* argue for a focus on drug plasma concentrations, in particular on active drug metabolites, as this has the potential to correlate to drug efficacy and is a defined quantitative phenotype [30]. Due to sample size, we focused on key candidate genes that were selected from large scale metabolome-wide association studies (MWAS) with the SNPs having a genetic associations for non-chlorinated endogenous cellular metabolites of the compounds of interest in our current study [27, 31, 32]. Two associations were identified in our study: first, a SLC7A5 SNP (rs28582913) was associated with 7-Cl-KYNA blood levels (p=0.00667). rs28582913 is an intronic variant with eQTL analysis showing a link to SLC7A5 gene expression in skin, tibial artery and adipose tissue (GTExPortal), with the CC allele having the lowest SLC7A5 gene expression. This correlates with our study where two CC alleles were associated with 46% lower blood concentrations. This suggests that LAT1 expression could be a rate limiting factor for the generation of the active metabolite as its expression determines (restricts) how much of the parent drug is able to cross at the BBB and thus have access to cells that express kynurenine aminotransferase (KATs) (Fig. 7). This is consistent with Lin *et al* who found that neural production of kynurenic acid requires AAT-1, a homologue of LAT1 [33]. However, a replication study would need to be carried out to build on our genetic association identified in the present study and to investigate if it correlates with drug response.

**Figure 7.**
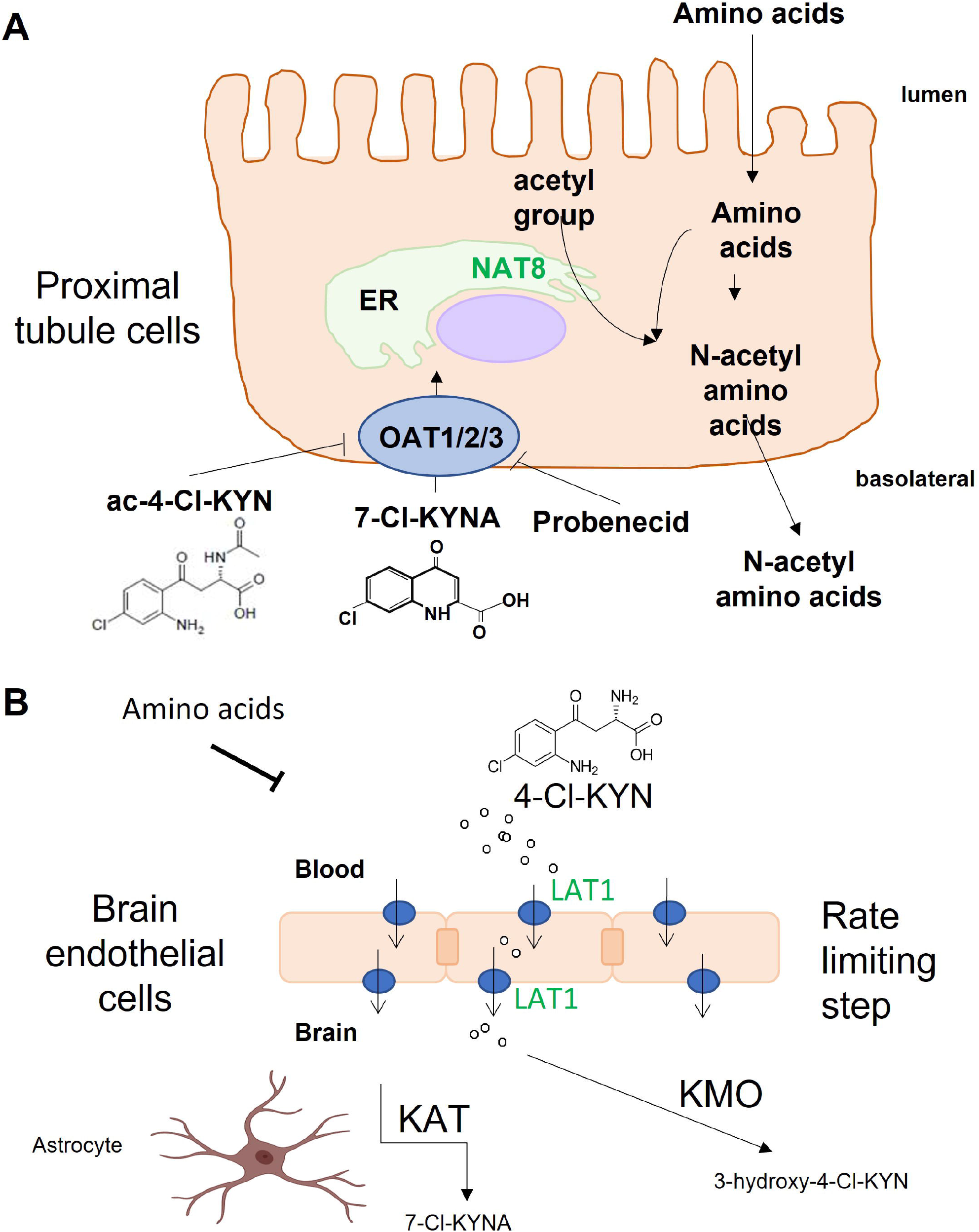
Passage of 4-CI-KYN metabolites, 7-CI-KYNA and N-acetyl-4-CI-KYN across renal proximal tubule cells and blood-brain barrier. **(A)** Diagram summarising the passage of the metabolites of prodrug, L-4-chlorokynurenine (4-CI-KYN) across renal proximal tubule cells. The active metabolite, 7-chlorokynurenic acid (7-CI-KYNA), is taken up into proximal tubule cells via organic anion transporters 1, 2 & 3 (OAT). Probenecid and the inactive metabolite of 4-CI-KYN (N­ acetyl-4-CI-KYN) inhibit OATs to prevent the uptake of 7-CI-KYNA across the basolateral membrane. Single nucleotide polymorphism in N-acetyltranferase 8 (NAT8) is linked to the ratio of N-acetyl-4-CI-KYN: 4-CI-KYN found in plasma. **(B)** Crossing of 4-CI-KYN across the blood-brain barrier by LAT1 (SLC7A5) and metabolism to active metabolite (7-CI-KYNA) in astrocytes or by KMO to 3-hydroxy-4-CI-KYN. SLC7A5 SNP is associated with the plasma level of 7-CI-KYNA.

Second, our data also revealed that the NAT8 SNP, rs13538, was associated with the ratio of N-acetyl-4-Cl-KYN: 4-Cl-KYN in plasma. NAT8 is predominantly expressed in the liver and kidneys and is involved in the removal of compounds by the addition of acetyl groups to amino acids or cystine conjugates to aid their excretion via bile or urine (Fig. 7). rs13538, a non-synonymous variant in NAT8, has been associated with circulating and urinary levels of n-acetyl-amino acids and has also been suggested as a susceptibility locus for chronic kidney disease [34, 35]. It has been speculated that higher blood concentrations of N-acetyl amino acids can be indicative of chronic kidney disease [35]. As well as rs13538 being a non-synonymous variant, eQTL analysis showed a link to NAT8 gene expression in tissues such as nerve and adipose tissue (GTExPortal) with the AA allele having the lowest NAT8 gene expression with an additive effect for each additional G allele at this variant. This correlates with our findings as patients with a GG allele at this locus had high acetyl-4-Cl-KYN:4-Cl-KYN ratio. As rs13538 is a non-synonymous variant, we investigated the location of this amino acid change using the AlphaFold [22] predictive structural model of NAT8. *In silico* docking studies with the putative substrate, 4-Cl-KYN, and acetyl Co-A found that they are in proximity within the binding pocket, in particular the transfer point of acetyl from donor to substrate is located near the Phe143Ser variant. The only functional studies for NAT8 involve work that identified the enzymatic activity of NAT8 to add an acetyl group to cysteine s-conjugates [36]. Therefore, it would be interesting to perform functional experiments to confirm the direct metabolism of amino acid to n-acetyl compounds as this has not been confirmed by enzyme-based metabolism assays and the involvement of the Phe143Ser variant.

In our study we have annotated our box and whisker plots of genetic associations with the self-identified ethnicity of the patients from the multicenter trial (ELEVATE) in the USA as our population cohort is 20% black. Underrepresented populations in pharmacogenetics are an active area of research as around 97% of GWAS data sets are of European ancestry while only 0.02% are of African American ethnicity [37, 38]. For our study, NAT8 ethnicity is known to alter the allele frequency; i.e. the rs13538 major allele A has a frequency of 0.79 in Europeans but the allele frequency is 0.38 in Africans. As the field of MWAS expands it will be interesting to investigate the relationship with ethnicity for the genes investigated but larger patient cohorts would be required for this work.

Due to the species differences in activity for specific substrates such as with kynurenic acid we investigated the transport of 7-Cl-KYNA by rodent and human OATs. An ongoing clinical trial (NCT03078322) will determine, if in humans, the co-administration of probenecid with 4-Cl-KYN will boost the CNS concentrations of 7-Cl-KYNA. OATs are proposed to have a role in sensing and signaling and so this would be an interesting avenue to investigate inter-organ communication following the co-administration of probenecid [39]. How probenecid co-administration would affect the genetic associations found in the current study is unknown and would require a cohort of patients co-administered with both 4-Cl-KYN and probenecid to study this.

In this study we used in vitro transporter assays, an animal study and a human data set to investigate the transport and metabolism of 4-Cl-KYN, as this increased knowledge base will enhance the development of this therapeutic agent (Fig. 7). For example, our finding of two genetic associations for drug concentration levels in plasma might enable a personalised medicine strategy for dosing, drug-drug interaction prediction and/or patient stratification for drug response. Further replication cohorts and studies are required for this.

## Acknowledgements

We thank Jo Cato, Senior Vice President of Development Operations, Vistagen Therapeutics, and Erik Berglund, Vice President of Global Regulatory Affairs, Vistagen Therapeutics for providing support to this research project.

The data used for the eQTL analyses described in this manuscript were obtained from the GTEx Portal on 30/09/22. The Genotype-Tissue Expression (GTEx) Project was supported by the Common Fund of the Office of the Director of the National Institutes of Health, and by NCI, NHGRI, NHLBI, NIDA, NIMH, and NINDS.

